# Retinoic acid signaling modulates smooth muscle cell phenotypic switching in atherosclerosis through epigenetic regulation of gene expression

**DOI:** 10.1101/2022.11.09.515888

**Authors:** Huize Pan, Sebastian E. Ho, Chenyi Xue, Jian Cui, Leila S. Ross, Muredach P. Reilly

## Abstract

**BACKGROUND:** Smooth muscle cells (SMCs) substantially contribute to the development of atherosclerosis through a process called “phenotypic switching.” Our previous work identified an SMC-derived intermediate cell type, termed “SEM” cells, which plays a crucial role in SMC transition to other cell types and in lesion development. Activation of retinoic acid (RA) signaling by all-trans retinoic acid (ATRA) attenuates atherosclerosis in mice coincident with suppression of SEM cell formation. However, the effect of RA signaling on advanced disease and the underlying molecular mechanisms by which RA modulates SMC transition to SEM cells are largely unknown.

**METHODS:** We applied SMC lineage tracing atheroprone mice and biochemistry and cell and molecular biology techniques (e.g., RNA sequencing, quantitative reverse transcription PCR, co-immunoprecipitation, and chromatin immunoprecipitation-quantitative PCR) to reveal the regulatory mechanisms of RA signaling in SMC transition to SEM cells.

**RESULTS:** Activation of RA signaling with ATRA in established atherosclerosis significantly reduced SEM cells and lesion size while increasing fibrous cap thickness. Mechanistically, retinoic acid receptor alpha (RARα) directly targets the promoters of *Ly6a* and *Ly6c1* in mouse SMCs, and activation of RA signaling recruits EZH2 to the regulatory elements triggering local H3K27me3. Distinct from a molecular model that reported for RA recruitment of HDAC1 during embryogenesis, RARα/EZH2 complex recruits SIRT1 and SIRT6, rather than classical HDACs, to the regulatory regions of key SEM cell marker genes. This subsequently reduces multiple acetylated histone modifications (e.g., H3K27ac, H3K4ac, H3K9ac, H3K14ac, H3K56ac) with recruitment of the transcription corepressor, NCOR1, to repress downstream SEM cell marker genes.

**CONCLUSIONS:** Our findings provide novel mechanistic insights into RA modulating SMC phenotypic switching in atherosclerosis, suggesting molecular targets for preventive and therapeutic interventions for atherosclerosis and its clinical complications.

## INTRODUCTION

Atherosclerosis, characterized by modulation and lesion infiltration of multiple cell types in response to lipid accumulation in the artery wall, remains the major cause of cardiovascular disease (CVD)^1^. Pathophysiological responses of multiple cell types, including vascular smooth muscle cells (SMCs), endothelial cells (ECs), and immunocytes (e.g., macrophages, T cells), contribute to the development, progression, and clinical complications of atherosclerosis^2^. SMCs contribute the majority of cells in atherosclerotic plaques through “phenotype switching” and transitioning to other cell types^3^. Several lines of evidence, including human genetics, SMC lineage tracing in mouse genetic models, and single cell RNA-sequencing (scRNA-seq)^4-7^, have shown that SMC phenotypic plasticity is a causal mediator for atherosclerotic plaque stability and clinical events, suggesting opportunities for therapeutic intervention for atherosclerotic CVD.

Our recent work and others identified multiple SMC-derived cell types in atherosclerotic lesions and several regulators of SMC phenotype switching, such as retinoic acid (RA) signaling, TCF21, KLF4, OCT4, ZEB2, in atherosclerosis^5-8^. Our previous research identified a novel cell state derived from SMCs, “SEM” cells, named because of their expression of marker genes of <u>s</u>tem cells (*Ly6a*), <u>e</u>ndothelial cells (*Vcam1*), and <u>m</u>onocytes (*Ly6c1*)^5^. SEM cells have substantial overlap with Lgals3^+^ intermediate cells^6^ and fibromyocytes^7^ described by other groups. We also indicated that activation of RA signaling with all-trans retinoic acid (ATRA) blocked SMC transition to SEM cells, attenuated atherosclerosis progression, and increased features of lesion stability^5^. Our current work seeks to understand the molecular mechanisms by which RA signaling modulates SMC-SEM cell transition.

Classically, RA (e.g., ATRA, 9-*cis*-RA, and 13-*cis*-RA) binds and activates RA receptors (RARs) to form heterodimers with retinoid X receptors (RXRs)^9, 10^. This complex binds to retinoic acid response elements (RAREs)^11^, which recruits histone acetyltransferase (HATs) and Trithorax proteins, mediating H3K4me3 and promoting downstream gene transcription^12^. However, some exceptions to this classical model have been reported. For instance, RA repression of development-related genes, such as *Hoxb1*^13^ and *Fgf8*^14^, was mediated by recruitment of EZH2 and HDAC1 to the upstream regulatory sequences of the genes. The regulatory mechanisms of RA signaling in SMC phenotype switching has yet to be elucidated.

Here, we demonstrate that even in advanced disease, activation of RA signaling by ATRA reduces the proportion and number of SEM cells and atherosclerotic lesion area while enhancing lesion stability. Using cultured mouse SMCs, we elucidated the precise molecular mechanism by which RA signaling represses expression of SEM cell marker genes and attenuates SMC-SEM cell transition. The mechanism involves RARα/EZH2 complex recruitment of SIRT1 and SIRT6, rather than classical histone deacetylases (HDACs), to the regulatory regions of SEM cell marker genes that represses their expression through extensive reduction in acetylated histone modifications and recruitment of the transcription corepressor, nuclear receptor corepressor 1 (NCOR1). Our study provides novel mechanistic insights into RA modulation of SMC phenotypic switching in atherosclerosis, suggesting molecular targets for clinical prevention and therapy for the disease.

## METHODS

### Mouse studies

All mouse experiments were approved by the Institutional Animal Care and Use Committee of Columbia University. Generation of SMC lineage tracing *ROSA26*^*LSL-ZsGreen1/+*^; *Ldlr*^−/−^; *Myh11-CreER*^*T2*^ mice and induction of SMC-specific expression of ZsGreen1 with tamoxifen diet and atherosclerosis with Western diet (WD) were previously described^5^. As the *Myh11*-CreER^T2^ locus was integrated into the Y chromosome, all SMC lineage tracing mice used in this study are male. To administer ATRA in advanced atherosclerosis, SMC lineage tracing mice were fed tamoxifen diet and then WD for 16 weeks before oral administration of ATRA (2.5 mg/kg mice) or vehicle (corn oil), 3 times/week.

### Hematoxylin and eosin staining

Apical parts of mouse hearts were collected, fixed, and embedded in Tissue Frozen Medium. Hematoxylin and eosin (H&E) staining was performed using aortic sinus sections in the Histology Service of the Molecular Pathology Shared Resource at the HICCC, Columbia University. Lesion area and fibrous cap thickness were measured using Aperio ImageScope software (Leica).

### Flow cytometry analysis

Single cells were prepared from mouse atherosclerotic aortas as previously reported^5^. Cell pellets were resuspended in FACS buffer with rat anti-mouse CD16/CD32 (eBioscience, 16-0161) to block unspecific binding of antibodies to Fc receptors and subsequently incubated with DAPI, rat anti-mouse LY6A-APC (eBioscience, 17-5981), and rat anti-mouse LY6C-PE (Biolegend, 128007) for 20 min at 4°C. SMC-derived ZsGreen1^+^LY6A^+^LY6C1^+^ SEM cells were analyzed and sorted on BD Influx instrument as previously described^5^.

### Cell culture and *in vitro* chemical treatment

Mouse smooth muscle cells and SEM cells were cultured in DMEM+10% FBS medium as previously described^5^. The cells were treated with vehicle, ATRA (10 μM), BMS493 (10 μM), and/or GSK126 (10 μM) as indicated for 3 days before analyses.

### RNA-seq data analyses

Paired-end strand-specific RNA libraries were sequenced on Illumina NovaSeq 6000. Gene expression was quantified using *Salmon* v0.11.2^15^ with transcriptome index built from GENCODE M20. Gene expression was obtained by summing transcript level TPM (Transcripts Per Million) for each gene. Scaled TPMs were used for visualization on heatmaps. Differential expression (DE) analysis was performed using *DESeq2* package in R^16^. Genes with ≥10 total count across all samples were retained for DE analysis. Genes with fold change ≥2, FDR <0.05 in the Wald test and FDR <0.05 in the likelihood ratio test were considered differentially expressed. Log2 fold change was shrunken using *apeglm* method^17^ implemented in DESeq2. Differentially expressed genes in ATRA group compared to control were subject to Gene Ontology (GO) enrichment analysis. Enriched Biological Pathways were identified for downregulated genes in ATRA group using *clusterProfiler* package^18^. GO terms were simplified using *simplify* function with a similarity cutoff of 0.7. Fastq files and gene expression matrix from RNA-seq data will be deposited in the Gene Expression Omnibus database. Accession code is pending.

### Quantitative RT-PCR

RNA was extracted and cDNA was synthesized according to the manufactures’ manuals of Quick-RNA Miniprep kit (Zymo Research, R1054) and High-Capacity cDNA Reverse Transcription Kit (Applied Biosystems, 4368813), respectively. Quantitative PCR (qPCR) was performed using 2x PowerUp SYBR Green Master Mix (Applied Biosystems, A25777) with QuantStudio 7 Flex Real-Time PCR System (Applied Biosystems). The following primers were used in qPCR experiments: mActb F: 5’-GGCTGTATTCCCCTCCATCG-3’; mActb R: 5’-CCAGTTGGTAACAATGCCATGT-3’; mLy6a F: 5’-AGGAGGCAGCAGTTATTGTGG-3’; mLy6a R: 5’-CGTTGACCTTAGTACCCAGGA-3’; mLy6c1 F: 5’-GCAGTGCTACGAGTGCTATGG-3’; mLy6c1 R: 5’-ACTGACGGGTCTTTAGTTTCCTT-3’.

### ChIP-qPCR

Chromatin immunoprecipitation quantitative PCR (ChIP-qPCR) was performed using the SimpleChIP Enzymatic Chromatin IP Kit (Magnetic Beads) (CST, #9003), following manufacturer’s protocol. The following antibodies were used for ChIP experiments: RARα (Abcam, ab41934), EZH2 (CST, #5246), Acetyl Histone H3 (Lys27) (CST, #8173), Tri-Methyl-Histone H3 (Lys27) (CST, #9733), SIRT1 (Millipore, 07-131), SIRT6 (Abcam, ab191385), Acetyl Histone H3 (Lys9) (Active Motif, 39137), Acetyl Histone H3 (Lys14) (Millipore, 07-353), Acetyl Histone H3 (Lys56) (Active Motif, 39281), Acetyl Histone H3 (Lys4) (Millipore, 07-539), NCOR1 (Bethyl Laboratories, A301-145A), p300 (Millipore, 05-257), Tri-Methyl-Histone H3 (Lys4) (CST, #9751), HDAC1 (CST, #34589), HDAC3 (CST, #85057), HDAC6 (CST, #7612), normal rabbit IgG (CST, #2729), and normal mouse IgG (Santa Cruz, sc-2025). The DNA products were quantified by qPCR using SimpleChIP Universal qPCR Master Mix (CST, #88989). The primers used for ChIP-qPCR are as follows: mLy6a promotor F: 5’-CTTCCATCCCAGTTGCCAGT-3’; mLy6a promotor R: 5’-CCCCAGTAGGCTCTTGCATC-3’; mLy6c1 promotor F: 5’-GCTTTCTAGTTGGCAAGCACA-3’; mLy6c1 promotor R: 5’-ACCATGGTGCAGGAGAACTG-3’.

### Co-IP and western blot

Co-immunoprecipitation (Co-IP) was conducted using the Nuclear Complex Co-IP Kit (Active Motif, 54001) according to the manufacturer’s manual. The antibodies used for Co-IP and following western blot are RARα (Abcam, ab41934), RARα (CST, #62294), EZH2 (CST, #5246), HDAC1 (CST, #34589), HDAC2 (CST, #57156), HDAC3 (CST, #85057), HDAC4 (CST, #15164), HDAC5 (CST, #20458), HDAC6 (CST, #7612), HDAC7 (CST, #33418), normal rabbit IgG (CST, #2729), and normal mouse IgG (Santa Cruz, sc-2025).

### Statistical analysis

Normality was assessed using the Shapiro-Wilk test. Statistical significance was determined using unpaired Student’s t-test for two group comparisons of atherosclerotic lesion areas of aortic sinuses, fibrous cap thickness, and SEM cell proportion and number from vehicle and ATRA-treated mice. RNA-seq, RT-qPCR, and ChIP-qPCR were performed in biological triplicates. Data are shown as mean±SD. GraphPad Prism 9 was used for all statistical analyses.

## RESULTS

We previously reported that ATRA attenuated SEM cell formation during atherosclerosis development and progression^5^. To examine the effects of ATRA on advanced disease, we fed *ROSA26*^*LSL-ZsGreen1/+*^; *Ldlr*^*-/-*^; *Myh11-CreER*^*T2*^ mice WD for 16 weeks, then gave another 10-week WD along with either ATRA or vehicle (corn oil) treatment (**Figure 1A**). ATRA treatment significantly reduced atherosclerotic lesion area and increased fibrous cap thickness in established atherosclerosis (**Figure S1A through S1C**). Importantly, among all ZsGreen1^+^ SMC lineage cells prepared from atherosclerotic aortas, we found that both the proportion and number of SEM cells were significantly decreased (**Figure 1B and 1C**).

**Figure 1.**
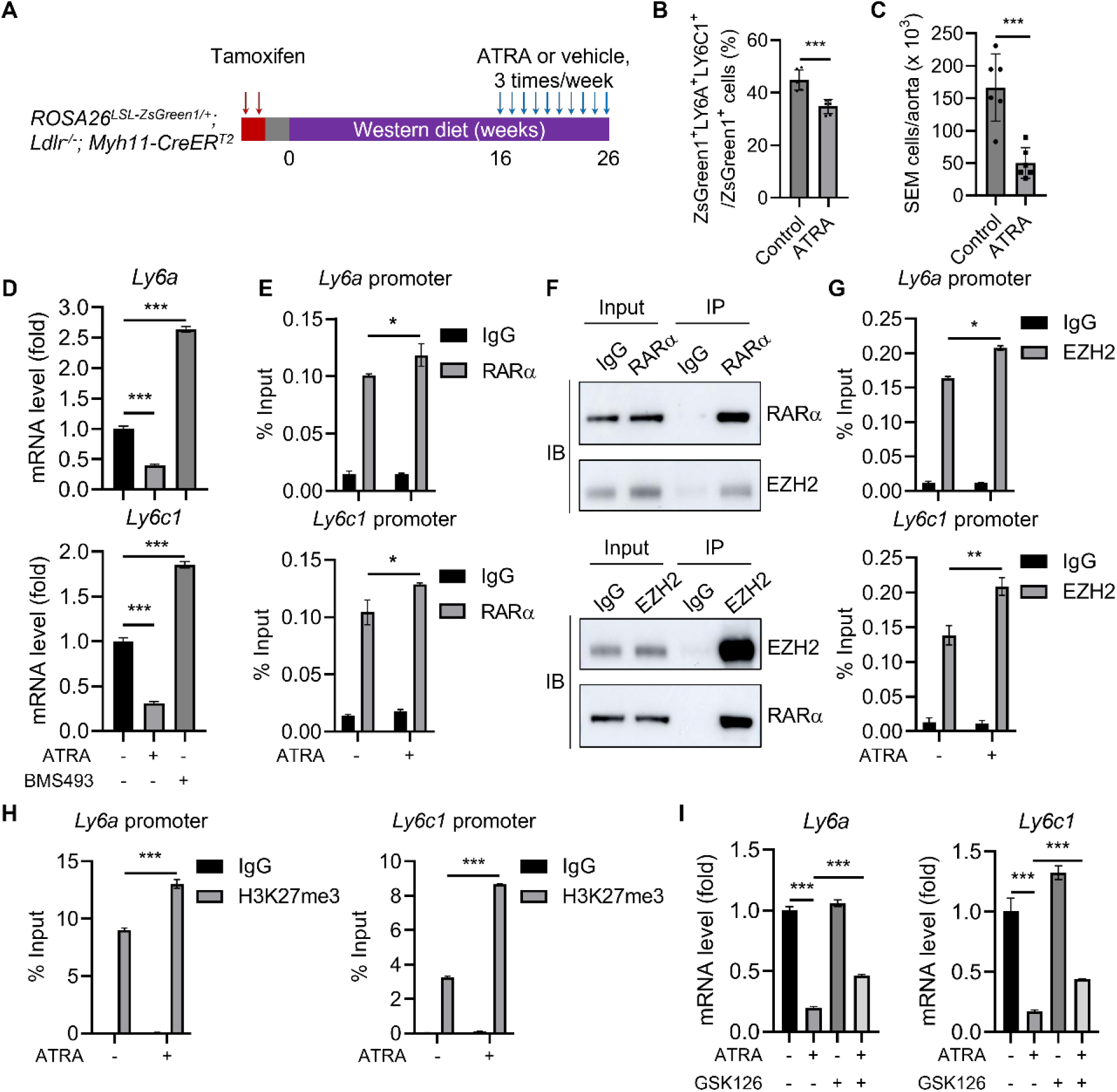
Activation of retinoic acid signaling represses SMC to SEM cell transition via EZH2-mediated epigenetic silencing of key SEM cell marker genes. **A**, Schematic of administration of ATRA and vehicle (corn oil), 3 times/week, to the SMC lineage tracing mice on Western diet for 26 weeks in total. **B** and **C**, Flow cytometry analysis of ZsGreen1^+^LY6A^+^LY6C1^+^ SEM cells showed both proportion (**B**) and number (**C**) of SEM cells from atherosclerotic aortas were markedly reduced by ATRA administration. n=6 mice for each group. \*\*\**P*<0.001. **D**, RT-qPCR of SEM cell marker genes, *Ly6a* and *Ly6c1*, in mouse SMCs indicated that activation of RA signaling by ATRA repressed while inhibition of RA signaling by BMS493 promoted SEM cell marker gene expression. n=3, \*\*\**P*<0.001. **E**, ChIP-qPCR showed enrichment of RARα at promoters of *Ly6a* and *Ly6c1*. n=3, \**P*<0.05. **F**, Endogenous co-IP with mouse SMC nuclear protein extractions indicated physical interaction between RARα and EZH2. **G**, ChIP-qPCR showed occupancy of EZH2 at promoters of *Ly6a* and *Ly6c1*, n=3, \**P*<0.05, \*\**P*<0.01. **H**, H3K27me3 levels at promoters of *Ly6a* and *Ly6c1* were significantly induced by ATRA treatment, n=3, \*\*\**P*<0.001. **I**, Treatment of mouse SMCs with GSK126, an EZH2 methyltransferase activity inhibitor, partially restored expression of SEM cell marker genes, *Ly6a* and *Ly6c1*, repressed by ATRA, n=3, \*\*\**P*<0.001.

To confirm that ATRA directly targets SMC to SEM cell transition through activation of RA signaling but not simply via changes in circulating lipoproteins as reported^19^, we treated *ex vivo* cultured SEM cells, pooled from atherosclerotic aortas, with ATRA. The expression of key SEM cell marker genes, *Ly6a* and *Ly6c1*, was markedly reduced by ATRA (**Figure S1D**). In addition to SEM cell marker genes, we previously observed that two other clusters of genes (i.e., extracellular matrix-related genes and inflammatory genes) were upregulated in SEM cells versus SMCs. Gene Ontology (GO) analysis of downregulated genes in SMCs treated with ATRA suggested that multiple biological processes, particularly “inflammatory response,” “extracellular matrix organization,” and “regulation of inflammatory response,” were repressed by ATRA (**Figure S2A**). Multiple genes related to extracellular matrix organization (e.g., *Fn1, Col1a2, Lum*) and inflammatory response (e.g., *Ccl7, Cxcl10, Ccl5*) were strongly downregulated in ATRA-treated SMCs (**Figure S2B through S2D**). These data, combined with our previous findings^5^, suggest that activation of RA signaling represses SMC transition to SEM cells by inhibiting expression of SEM cell-related genes, particularly the key SEM cell marker genes.

To further corroborate that SEM cell marker genes are directly repressed by RA signaling, rather than non-RAR target effects of ATRA, we treated mouse SMCs with ATRA or BMS493, an inverse agonist of pan-RARs. Inhibition of RA signaling by BMS493 significantly upregulated SEM cell marker genes, *Ly6a* and *Ly6c1* (**Figure 1D**). To test if RA directly regulates transcription of SEM cell marker genes, we examined enrichment of RARα, a key receptor for RA ligands, at promoter regions of *Ly6a* and *Ly6c1*. RARα occupied both *Ly6a* and *Ly6c1* promoters at baseline, which was induced further by ATRA treatment (**Figure 1E)**. These data indicate that RA signaling can directly target transcription of key SEM cell marker genes.

Activation of RA signaling typically replaces transcriptional repressors with co-activators, which then recruit HATs, activating downstream gene transcription^12^. However, we did not find enrichment of p300, a HAT reported to participate in RA-mediated gene activation^20^, at the promoters of *Ly6a* and *Ly6c1* under basal or ATRA-treated conditions (**Figure S3A**). Therefore, we tested an alternative mechanism that has been reported for RA-induced gene repression in development, involving recruitment of EZH2 to trigger H3K27me3^14^. First, our endogenous co-immunoprecipitation (co-IP) results in mouse SMCs indicated a physical interaction between RARα and EZH2 (**Figure 1F**). Next, we found that EZH2 directly targeted the promoter regions of *Ly6a* and *Ly6c1* and enrichment of EZH2 at the promoters increased in response to activation of RA signaling by ATRA (**Figure 1G)**.

EZH2 is a histone methyltransferase known to induce H3K27me3 at promoters to repress downstream genes^21^. We found that ATRA induced a marked upregulation of H3K27me3 at *Ly6a* and *Ly6c1* promoters in mouse SMCs (**Figure 1H)**. In contrast, the levels of H3K4me3, another active histone methylation mark, at the promoters of *Ly6a* and *Ly6c1* were comparable in ATRA versus vehicle-treated SMCs (**Figure S3B**). Moreover, GSK126, a highly selective EZH2 methyltransferase inhibitor^22^, partially rescued the ATRA-repressed *Ly6a* and *Ly6c1* expression (**Figure 1I**), suggesting that EZH2 methyltransferase activity is necessary for the suppression of SEM cell marker genes. Overall, these data suggest a model of RA-mediated recruitment of EZH2 triggering H3K27me3 at promoters of SEM cell marker genes to suppress their transcription.

Other transcription cofactors, such as HDACs (e.g., HDAC1), are reported to participate in RARα/EZH2-mediated gene repression in development (e.g., *Hoxb1, Fgf8*)^13, 14^ through modulation of histone acetylation at promoters of downstream genes. However, it is unclear if HDACs participate in regulation of SEM cell marker genes. We found that levels of H2K27ac and H3K4ac, two types of histone acetylation related to active gene transcription, were decreased at promoters of *Ly6a* and *Ly6c1* in response to activation of RA signaling in SMCs (**Figure S3C and S3D**). Mammalian HDACs can be divided into several classes, such as Class I (HDAC1-3), Class II (HDAC4-7), and Class III (SIRT1-7)^23^. Our co-IP data suggested that RARα physically interacted with HDAC1, HDAC3, and HDAC6 in SMCs (**Figure S4A**). However, ChIP-qPCR revealed decreased occupancy of HDAC1 and HDAC3 (**Figure S4B and S4C**) and no significant enrichment of HDAC6 (**Figure S4D**) at the promoters of *Ly6a* and *Ly6c1* in response to ATRA. This suggests that Class I and Class II HDACs may not be responsible for the decreased levels of H3K27ac and H3K4ac at promoter regions of SEM cell marker genes.

Next, we focused on SIRT1 and SIRT6 as they are deacetylases predominantly found in the nucleus and regulate gene expression^24^. We found that SIRT1 and SIRT6 occupied the promoters of *Ly6a* and *Ly6c1* and that their enrichment at the promoters was induced by ATRA (**Figure 2A and 2B**), suggesting that SIRT1 and SIRT6 participate in RA-mediated SEM cell marker gene repression by catalyzing several histone acetylation modifications at the promoters of *Ly6a* and *Ly6c1*. To further address this, we examined additional known substrates of SIRT1 and SIRT6 deacetylases (H3K9ac, H3K14ac, and H3K56ac; epigenetic marks for active gene transcription)^25^ and found that occupancy of all three epigenetic modifications of histone acetylation was decreased at promoters of *Ly6a* and *Ly6c1* by ATRA (**Figure 2C through 2E**).

**Figure 2.**
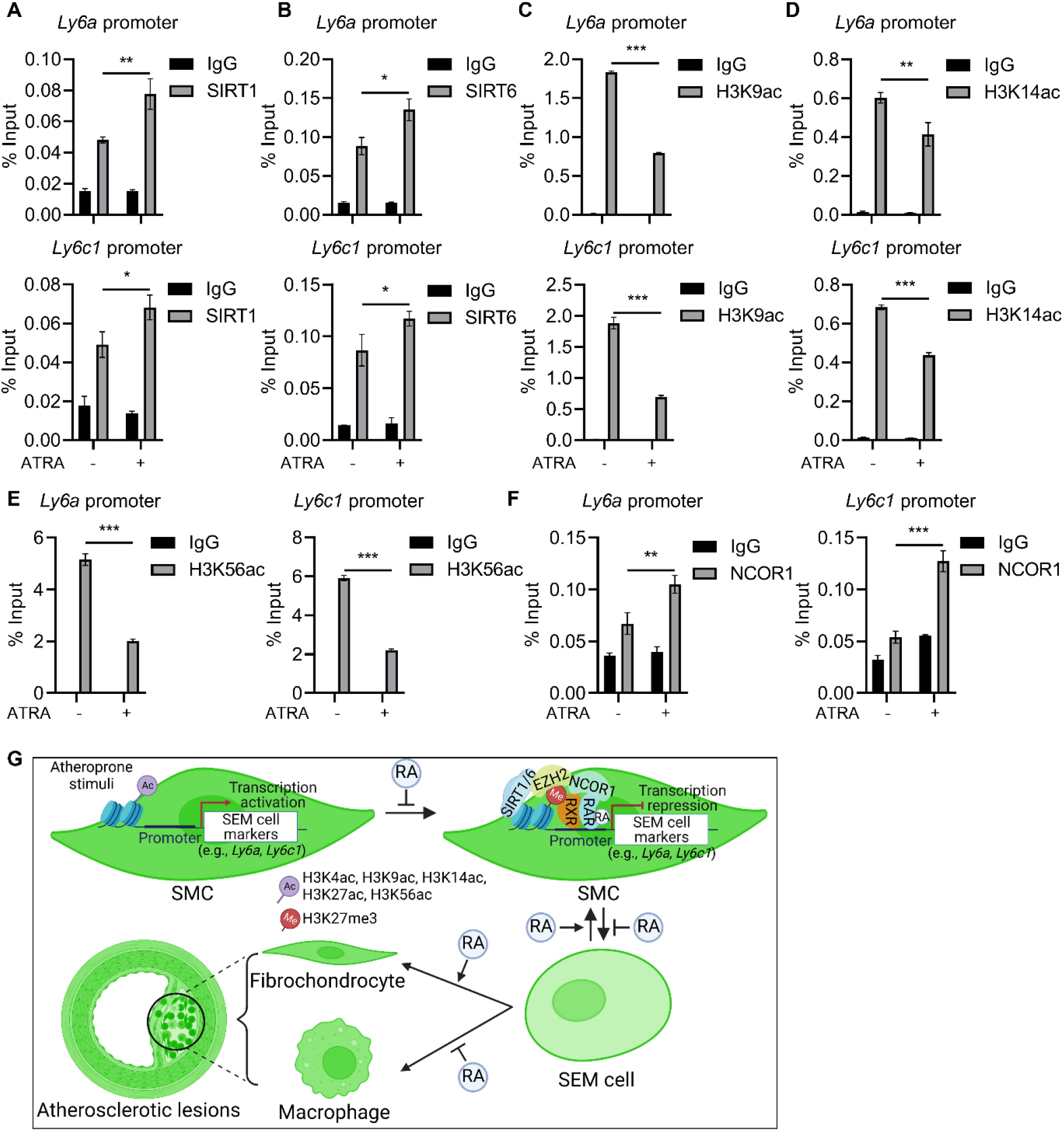
SIRT1 and SIRT6 mediate RARα/EZH2 repression of SEM cell marker genes through extensive alterations in epigenetic marks at promoter regions. **A** and **B**, ChIP-qPCR results showed that enrichment of SIRT1 and SIRT6 at promoters of *Ly6a* and *Ly6c1* was enhanced in response to activation of RA signaling by ATRA. n=3, \**P*<0.05, \*\**P*<0.01. **C** through **E**, ChIP-qPCR data show that levels of SIRT1 and SIRT6 substrates, H3K9ac (**C**), H3K14ac (**D**), and H3K56ac (**E**), at promoter regions of SEM cell marker genes, *Ly6a* and *Ly6c1*, were significantly reduced by ATRA. n=3, \*\**P*<0.01, \*\*\**P*<0.001. **F**, Transcription corepressor, NCOR1, was enriched at promoters of *Ly6a* and *Ly6c1* by activation of RA signaling. **G**, Schematic of proposed molecular model by which RA signaling modulates SMC to SEM cell transition in atherosclerosis and affects disease development.

Alterations of these epigenetic modifications (H3K27me3, H3K27ac, H3K4ac, H3K9ac, H3K14ac, and H3K56ac) result in chromatin compaction and recruitment of transcription corepressors to block downstream gene transcription^26^. To consolidate this, we interrogated the enrichment of NCOR1, a transcription corepressor associated with RA signaling in autophagy modulation^27^, at regulatory elements of SEM cell marker genes. The results indicated that ATRA led to markedly increased recruitment of NCOR1 to the promoters of *Ly6a* and *Ly6c1* (**Figure 2F**), suggesting that RA-altered epigenetic modifications at promoters of SEM cell marker genes inhibits downstream genes at least partially through recruitment of NCOR1.

## DISCUSSION

SMC phenotype switching is a central event during development of atherosclerosis. Several signaling pathways (e.g., RA signaling^5^), transcription factors (e.g., TCF21^7^, KLF4, OCT4^6^), and epigenetic modifiers (e.g., ZEB2^8^) have been identified as regulators of SMC phenotype switching and their clinically importance is supported by human genetic studies. However, the molecular mechanisms of how these regulators modulate SMC transition to SEM cells are only partially understood. The effects of RA signaling on SMC proliferation^28^ and neointima formation in a rat injury model^29^ were explored two decades ago. In subsequent studies, activation of RA signaling was found to decrease atherosclerotic lesion area in rodent atheroprone models^19, 30^. Yet until our recent work^5^, the actions of RA signaling on SMC phenotype switching and trajectories of SMC-derived cells in atherosclerosis were not known. Here, we focused on the molecular regulatory mechanisms of RA signaling in SMC phenotype switching in atherosclerosis.

First, we found that activation of RA signaling can suppress SEM cells and regress atherosclerotic lesions in mice with advanced disease, providing a translational framework for therapeutic targeting of established disease not just prevention of atherosclerosis as suggested by our prior studies^5^. Next, we demonstrated a novel molecular model for how RA signaling suppresses expression of key SEM cell marker genes (e.g., *Ly6a* and *Ly6c1*) and SMC transition to SEM cells. Activation of RA signaling recruited EZH2 to the promoters of key SEM cell marker genes, triggering the repressive epigenetic mark, H3K27me3, and enhanced SIRT1 and SIRT6 deacetylases, reducing histone acetylation at multiple lysine residues, marks of active gene transcription, at promoters of *Ly6a* and *Ly6c1*. The relevant chromatin remodeling recruits the transcription corepressor, NCOR1, to repress expression of downstream SEM cell marker genes. By doing so, RA signaling inhibits SMC to SEM cell transition and attenuates atherosclerosis (**Figure 2G**).

The novel molecular model of RA-mediated suppression of SMC to SEM cell transition that we revealed is distinct from the classical RA-induced gene activation paradigm. Similar to a few prior reports of RA repression of genes (e.g., *Hoxb1*^13^, *Fgf8*^14^) during mouse embryogenesis, we found that RA signaling blocked expression of SEM cell marker genes, *Ly6a* and *Ly6c1*, by recruiting EZH2 and triggering H3K27me3 at the promoters of the target genes. Yet, inconsistent with these prior reports, we did not find subsequent recruitment of HDAC1 or other classical HDACs (e.g., HDAC2-7) by RARα/EZH2 complex. Instead, SIRT1 and SIRT6 were markedly enriched at *Ly6a* and *Ly6c1* promoters in response to the activation of RA signaling, resulting in decrease of multiple acetylated histone modifications (e.g., H3K27ac, H3K4ac, H3K9ac, H3K14ac, H3k56ac) at these promoter regions. Our current work is the first to reveal that SIRT1 and SIRT6 are transcription cofactors participating in RA signaling-mediated suppression of SEM cell marker genes and blocking of SMC transition to SEM cells.

Our research provides novel insights but has limitations that merit further work. We have identified RARα/EZH2/SIRT1/SIRT6/NCOR1 complex in RA repression of SMC to SEM cells, but this does not exclude the potential involvement of additional transcriptional cofactors (e.g., HDAC8-11, HATs, or histone methyltransferases). Our data indicate that RA modulates gene expression in a broad epigenetic regulatory manner to block SMC to SEM cell transition. Indeed, it has been widely reported that multiple epigenetic remodelers, such as HATs (e.g., NCOA3^31^), HDACs (e.g., HDAC1^14^), are involved in RA-mediated gene regulation in other settings. Future studies are needed to interrogate the broader epigenetic landscape alterations that are driven by activation of RA signaling and how these are deterministic of SMC transition to SEM cells and subsequent transition to other cell types.

There are significant differences and overlap in the atherosclerosis phenotypes described for each of the signaling (e.g., RA signaling), genetic (e.g., TCF21, KLF4), and epigenetic (e.g., ZEB2) regulators of SMC phenotype switching. For instance, *Klf4* deletion in SMCs results in suppression of SMC phenotypic transition and marked reduction in atherosclerotic lesion size^6^. SMC-specific genetic deletion of *Tcf21* leads to decreased SMC-derived fibromyocytes within lesions and fibrous caps^7^. Loss of *Zeb2* in SMCs inhibits SMC transition to fibromyocytes but promotes chondromyocyte development^8^. In contrast, activation of RA signaling by ATRA blocks SMC to SEM cell transition, attenuates atherosclerosis, and promotes lesion stability^5^. Given this phenotypic overlap, there may be crosstalk in the regulatory events among these factors and signaling pathways in SMCs. Indeed, it has been reported that RA induced *Klf4* expression in SMCs^32^. Our data provide the context to interrogate, at the local and genome level, specific interactions between RA signaling and other regulators of SMC phenotypic switching.

In conclusion, we reveal a novel molecular mechanism by which RA signaling blocks SMC to SEM cell transition, providing a deeper mechanistic insight into SMC phenotypic switching in atherosclerosis. Our study suggests several molecular targets for both prevention of atherosclerosis progression and therapy of established disease independent of lipid lowering strategies currently used in clinic.

## Acknowledgments

Block sectioning and H&E staining were performed in the Columbia Molecular Pathology Shared Resource of HICCC, supported by NIH/NCI grant #P30CA013696. Flow cytometry analysis and FACS sorting were performed in the Columbia CCTI Flow Cytometry Core, supported in part by the NIH award S10OD020056. The RNA-seq was performed in JP Sulzberger Columbia Genome Center, funded in part through the NIH/NCI Cancer Center Support Grant P30CA013696.

## Sources of Funding

This work is supported by NIH K99/R00 Award (K99HL153939, H.P.). M.P.R. is supported by NIH grants R01HL113147 and R01HL150359.

## Disclosures

None.

## Supplementary Figures

**Figure S1.**
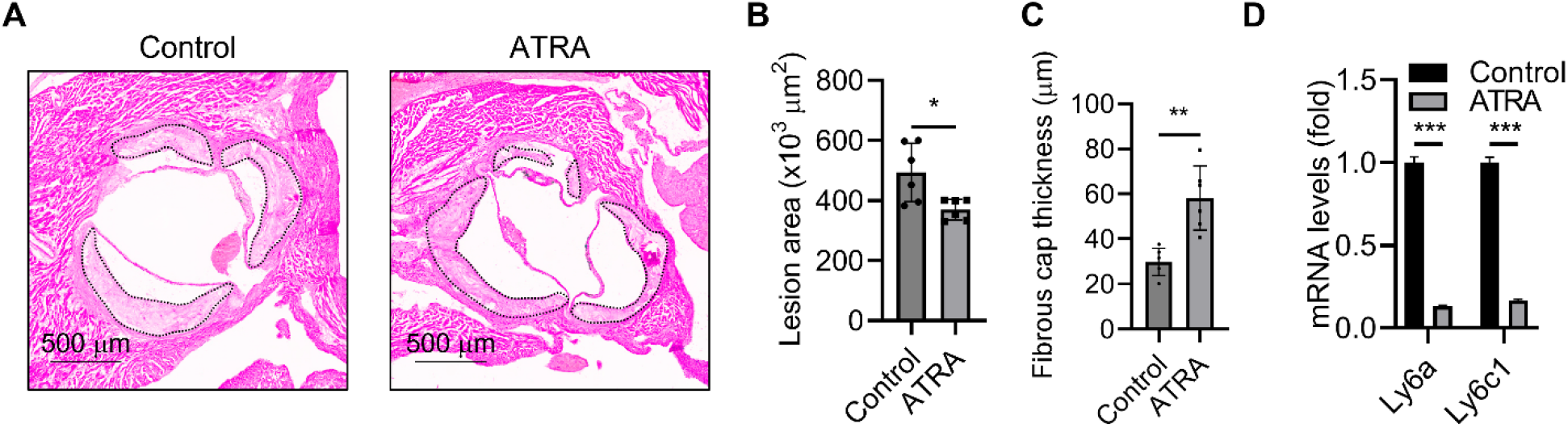
Activation of retinoic acid signaling in advanced atherosclerosis blocks SMC-derived SEM cells, reduces lesion area, and increases fibrous cap thickness. **A**, Representative images of hematoxylin and eosin (H&E) stained aortic root sections from vehicle (Control) and ATRA treated mice on Western diet for 26 weeks. **B** and **C**, Statistical analysis of lesion areas (**B**) and fibrous cap thickness (**C**) of aortic root sections from mice in vehicle (Control) and ATRA groups. n=6 mice for each group. \**P*<0.05, \*\**P*<0.01. **D**, Expression of SEM cell marker genes, *Ly6a* and *Ly6c1*, in *ex vivo* cultured SEM cells was dramatically reduced by ATRA treatment. n=3, \*\*\**P*<0.001.

**Figure S2.**
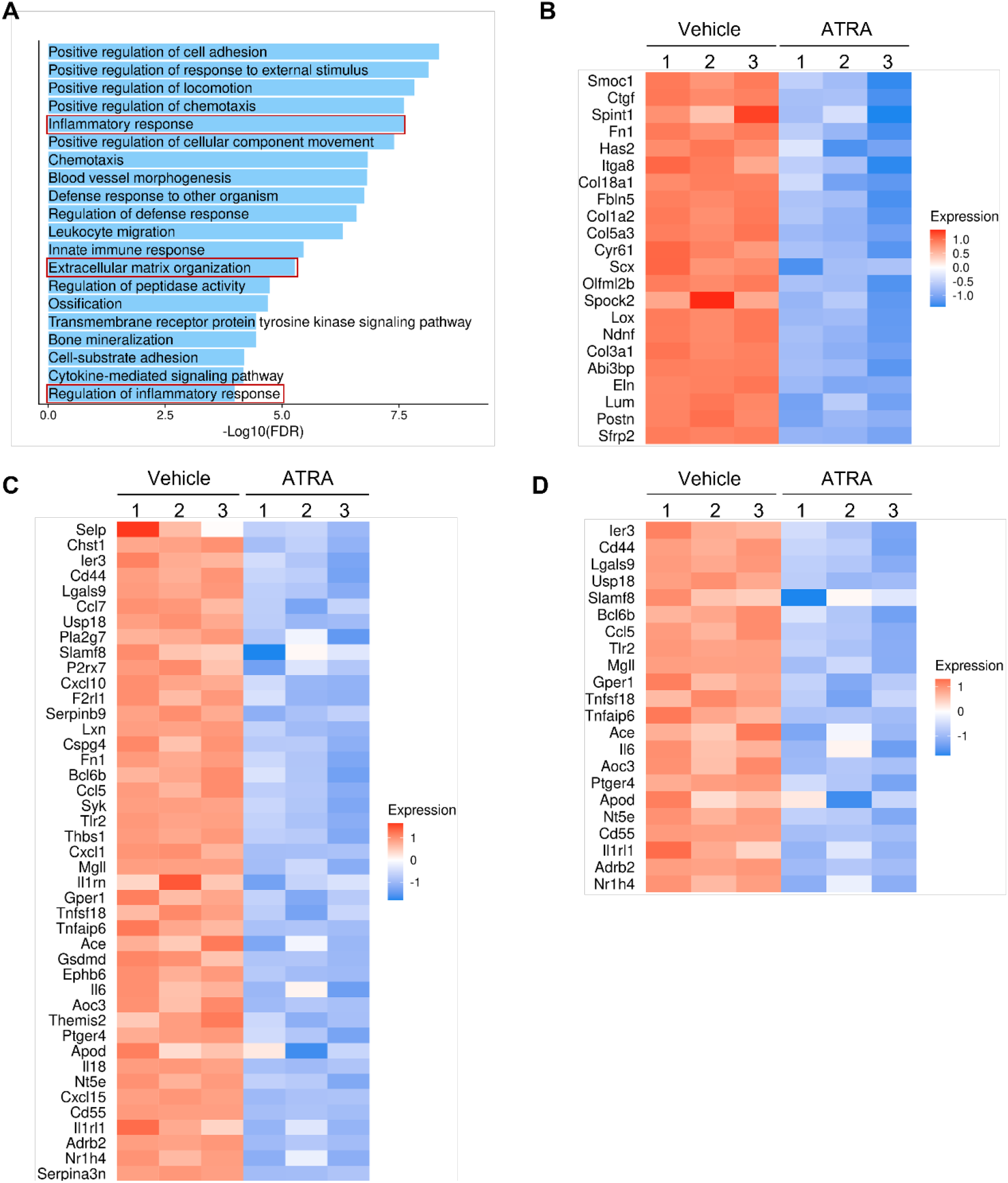
Activation of retinoic acid signaling represses multiple SMC-derived SEM cell related genes and pathways. **A**, Gene Ontology (GO) analysis of downregulated genes in ATRA-treated mouse SMCs versus vehicle group indicated multiple biological processes, including “inflammatory response,” “extracellular matrix organization,” and “regulation of inflammatory response,” were repressed by activation of retinoic acid signaling. **B** through **D**, heatmaps showing that multiple SEM cell-associated genes, including extracellular matrix related genes (**B**) and inflammatory genes (**C** and **D**), were markedly suppressed by ATRA.

**Figure S3.**
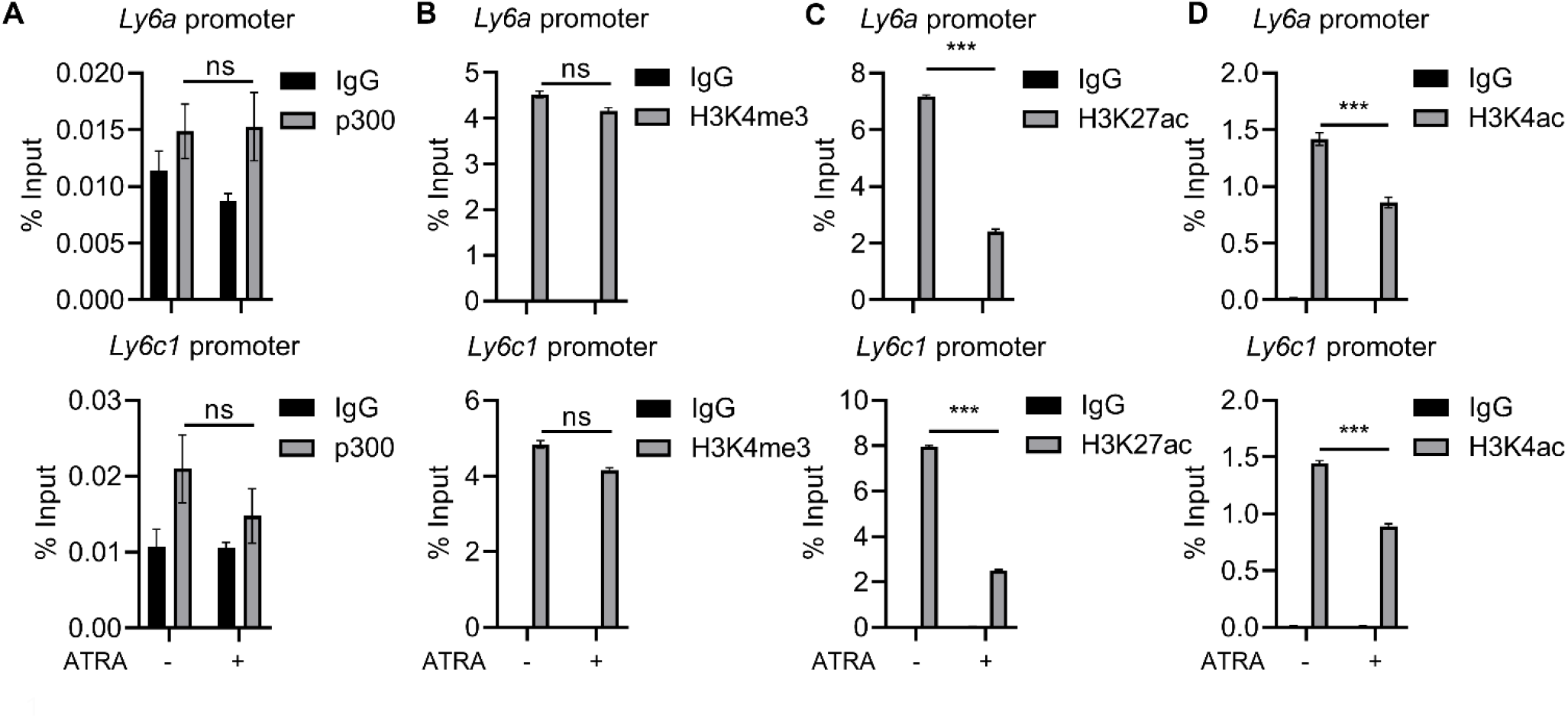
Enrichment of p300, H3K4me3, H3K27ac, and H3K4ac at promoters of key SEM cell marker genes in vehicle and ATRA treated mouse SMCs. **A** and **B**, ChIP-qPCR showed comparable enrichment of p300 and epigenetic mark H3K4me3 at promoters of SEM cell marker genes, *Ly6a* and *Ly6c1*, in vehicle and ATRA treated mouse SMCs. ns, not significant. **C** and **D**, Levels of H3K27ac and H3K4ac at promoters of *Ly6a* and *Ly6c1* were significantly induced by ATRA. n=3, \*\*\**P*<0.001.

**Figure S4.**
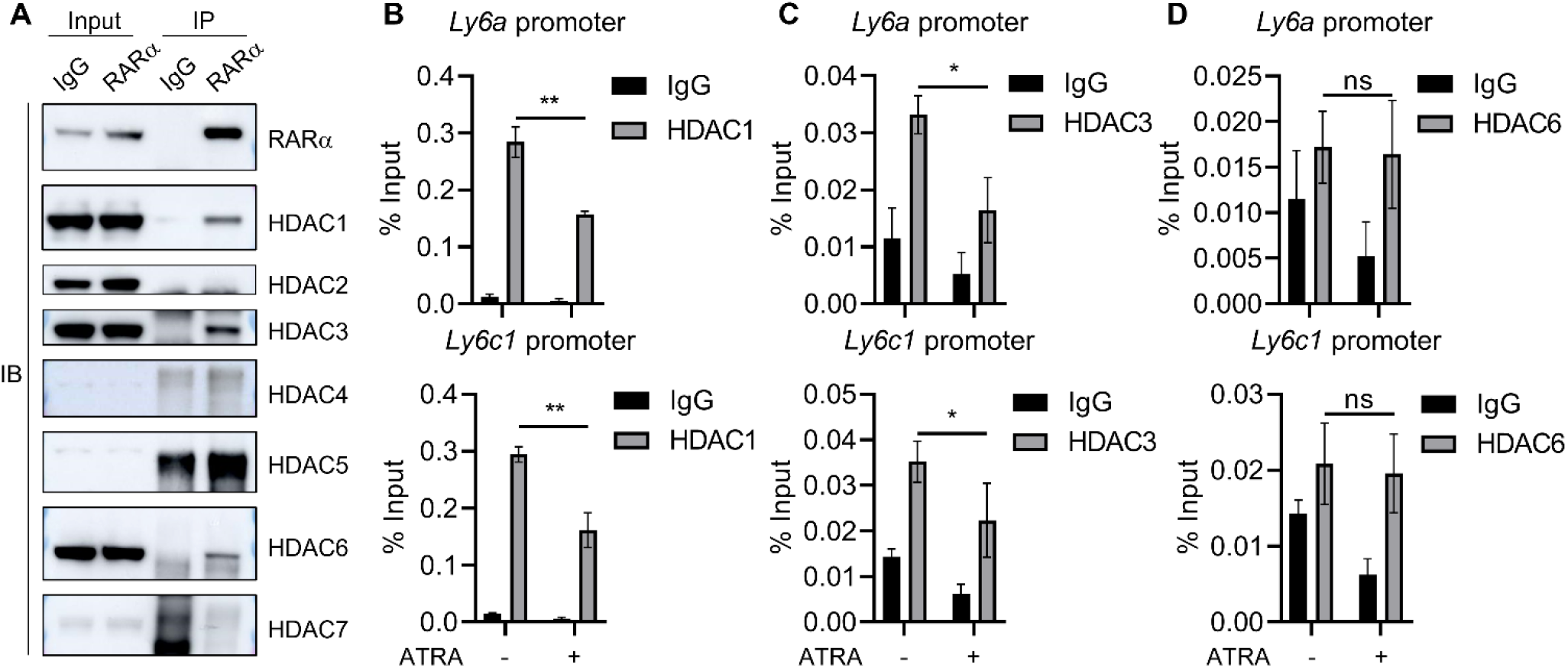
HDAC1-7 do not participate in RARα/EZH2-mediated SEM cell maker gene repression. **A**, Endogenous co-IP with mouse SMC nuclear protein extractions indicated physical interaction between RARα and HDAC1, HDAC3 and HDAC6. Note: IgG heavy chain bands showed in exposure images of HDAC4, HDAC5, and HDAC7. **B** and **C**, ChIP-qPCR showing reduced enrichment of HDAC1 and HDAC3 at promoters of *Ly6a* and *Ly6c1* in ATRA-treated mouse SMCs. n=3, \**P*<0.05, \*\**P*<0.01. **D**, ChIP-qPCR data indicated little occupancy of HDAC7 at promoters of *Ly6a* and *Ly6c1* in both vehicle and ATRA-treated mouse SMCs. ns, not significant.

## Notes

### Competing Interest Statement

The authors have declared no competing interest.

## REFERENCES

1. Herrington W, Lacey B, Sherliker P, Armitage J and Lewington S. Epidemiology of Atherosclerosis and the Potential to Reduce the Global Burden of Atherothrombotic Disease. Circ Res. 2016;118:535–46.

2. Stary HC, Chandler AB, Dinsmore RE, Fuster V, Glagov S, Insull W, Jr., Rosenfeld ME, Schwartz CJ, Wagner WD and Wissler RW. A definition of advanced types of atherosclerotic lesions and a histological classification of atherosclerosis. A report from the Committee on Vascular Lesions of the Council on Arteriosclerosis, American Heart Association. Arterioscler Thromb Vasc Biol. 1995;15:1512–31.

3. Basatemur GL, Jorgensen HF, Clarke MCH, Bennett MR and Mallat Z. Vascular smooth muscle cells in atherosclerosis. Nat Rev Cardiol. 2019;16:727–744.

4. Gomez D and Owens GK. Smooth muscle cell phenotypic switching in atherosclerosis. Cardiovasc Res. 2012;95:156–64.

5. Pan H, Xue C, Auerbach BJ, Fan J, Bashore AC, Cui J, Yang DY, Trignano SB, Liu W, Shi J, Ihuegbu CO, Bush EC, Worley J, Vlahos L, Laise P, Solomon RA, Connolly ES, Califano A, Sims PA, Zhang H, Li M and Reilly MP. Single-Cell Genomics Reveals a Novel Cell State During Smooth Muscle Cell Phenotypic Switching and Potential Therapeutic Targets for Atherosclerosis in Mouse and Human. Circulation. 2020;142:2060–2075.

6. Alencar GF, Owsiany KM, Karnewar S, Sukhavasi K, Mocci G, Nguyen AT, Williams CM, Shamsuzzaman S, Mokry M, Henderson CA, Haskins R, Baylis RA, Finn AV, McNamara CA, Zunder ER, Venkata V, Pasterkamp G, Bjorkegren J, Bekiranov S and Owens GK. Stem Cell Pluripotency Genes Klf4 and Oct4 Regulate Complex SMC Phenotypic Changes Critical in Late-Stage Atherosclerotic Lesion Pathogenesis. Circulation. 2020;142:2045–2059.

7. Wirka RC, Wagh D, Paik DT, Pjanic M, Nguyen T, Miller CL, Kundu R, Nagao M, Coller J, Koyano TK, Fong R, Woo YJ, Liu B, Montgomery SB, Wu JC, Zhu K, Chang R, Alamprese M, Tallquist MD, Kim JB and Quertermous T. Atheroprotective roles of smooth muscle cell phenotypic modulation and the TCF21 disease gene as revealed by single-cell analysis. Nat Med. 2019;25:1280–1289.

8. Cheng P, Wirka RC, Shoa Clarke L, Zhao Q, Kundu R, Nguyen T, Nair S, Sharma D, Kim HJ, Shi H, Assimes T, Brian Kim J, Kundaje A and Quertermous T. ZEB2 Shapes the Epigenetic Landscape of Atherosclerosis. Circulation. 2022;145:469–485.

9. Petkovich M, Brand NJ, Krust A and Chambon P. A human retinoic acid receptor which belongs to the family of nuclear receptors. Nature. 1987;330:444–50.

10. Giguere V, Ong ES, Segui P and Evans RM. Identification of a receptor for the morphogen retinoic acid. Nature. 1987;330:624–9.

11. Niederreither K and Dolle P. Retinoic acid in development: towards an integrated view. Nat Rev Genet. 2008;9:541–53.

12. Cunningham TJ and Duester G. Mechanisms of retinoic acid signalling and its roles in organ and limb development. Nat Rev Mol Cell Biol. 2015;16:110–23.

13. Studer M, Popperl H, Marshall H, Kuroiwa A and Krumlauf R. Role of a conserved retinoic acid response element in rhombomere restriction of Hoxb-1. Science. 1994;265:1728–32.

14. Kumar S and Duester G. Retinoic acid controls body axis extension by directly repressing Fgf8 transcription. Development. 2014;141:2972–7.

15. Patro R, Duggal G, Love MI, Irizarry RA and Kingsford C. Salmon provides fast and bias-aware quantification of transcript expression. Nat Methods. 2017;14:417–419.

16. Soneson C, Love MI and Robinson MD. Differential analyses for RNA-seq: transcript-level estimates improve gene-level inferences. F1000Res. 2015;4:1521.

17. Zhu A, Ibrahim JG and Love MI. Heavy-tailed prior distributions for sequence count data: removing the noise and preserving large differences. Bioinformatics. 2019;35:2084–2092.

18. Yu G, Wang LG, Han Y and He QY. clusterProfiler: an R package for comparing biological themes among gene clusters. OMICS. 2012;16:284–7.

19. Zhou B, Pan Y, Hu Z, Wang X, Han J, Zhou Q, Zhai Z and Wang Y. All-trans-retinoic acid ameliorated high fat diet-induced atherosclerosis in rabbits by inhibiting platelet activation and inflammation. J Biomed Biotechnol. 2012;2012:259693.

20. Kawasaki H, Eckner R, Yao TP, Taira K, Chiu R, Livingston DM and Yokoyama KK. Distinct roles of the co-activators p300 and CBP in retinoic-acid-induced F9-cell differentiation. Nature. 1998;393:284–9.

21. Cao R and Zhang Y. The functions of E(Z)/EZH2-mediated methylation of lysine 27 in histone H3. Curr Opin Genet Dev. 2004;14:155–64.

22. McCabe MT, Ott HM, Ganji G, Korenchuk S, Thompson C, Van Aller GS, Liu Y, Graves AP, Della Pietra A, 3rd, Diaz E, LaFrance LV, Mellinger M, Duquenne C, Tian X, Kruger RG, McHugh CF, Brandt M, Miller WH, Dhanak D, Verma SK, Tummino PJ and Creasy CL. EZH2 inhibition as a therapeutic strategy for lymphoma with EZH2-activating mutations. Nature. 2012;492:108–12.

23. Seto E and Yoshida M. Erasers of histone acetylation: the histone deacetylase enzymes. Cold Spring Harb Perspect Biol. 2014;6:a018713.

24. Kalous KS, Wynia-Smith SL, Summers SB and Smith BC. Human sirtuins are differentially sensitive to inhibition by nitrosating agents and other cysteine oxidants. J Biol Chem. 2020;295:8524–8536.

25. Gamez-Garcia A and Vazquez BN. Nuclear Sirtuins and the Aging of the Immune System. Genes (Basel). 2021;12.

26. Struhl K. Histone acetylation and transcriptional regulatory mechanisms. Genes Dev. 1998;12:599–606.

27. Gomez-Sintes R, Xin Q, Jimenez-Loygorri JI, McCabe M, Diaz A, Garner TP, Cotto-Rios XM, Wu Y, Dong S, Reynolds CA, Patel B, de la Villa P, Macian F, Boya P, Gavathiotis E and Cuervo AM. Targeting retinoic acid receptor alpha-corepressor interaction activates chaperone-mediated autophagy and protects against retinal degeneration. Nat Commun. 2022;13:4220.

28. Miano JM, Topouzis S, Majesky MW and Olson EN. Retinoid receptor expression and all-trans retinoic acid-mediated growth inhibition in vascular smooth muscle cells. Circulation. 1996;93:1886–95.

29. Miano JM, Kelly LA, Artacho CA, Nuckolls TA, Piantedosi R and Blaner WS. all-Trans-retinoic acid reduces neointimal formation and promotes favorable geometric remodeling of the rat carotid artery after balloon withdrawal injury. Circulation. 1998;98:1219–27.

30. Zhou W, Lin J, Chen H, Wang J, Liu Y and Xia M. Retinoic acid induces macrophage cholesterol efflux and inhibits atherosclerotic plaque formation in apoE-deficient mice. Br J Nutr. 2015;114:509–18.

31. Kashyap V and Gudas LJ. Epigenetic regulatory mechanisms distinguish retinoic acid-mediated transcriptional responses in stem cells and fibroblasts. J Biol Chem. 2010;285:14534–48.

32. Shi JH, Zheng B, Chen S, Ma GY and Wen JK. Retinoic acid receptor alpha mediates all-trans-retinoic acid-induced Klf4 gene expression by regulating Klf4 promoter activity in vascular smooth muscle cells. J Biol Chem. 2012;287:10799–811.

